# A Hybridization-Based Novel DNA Storage Technology with Rapid Read Performance

**DOI:** 10.64898/2025.12.24.696143

**Authors:** Yifei Zhang, Lijia Jia, Xiaoxia Wang, Yue Shi, Haihua Wang, Dongyi Zhang, Pengfei Cheng, Ye Zhao, Mingyuan Xu, Xinyi Zhang, Zuying Xiao, Lei Wu, Xianghong Zhou, Di Liu

## Abstract

The development of traditional silicon-based microelectronic storage media has increasingly lagged behind the rapidly growing demands of contemporary data storage. DNA storage, with its ultra-high information density and long-term preservation advantages, has rapidly become a research hotspot for next-generation storage technologies. However, current mainstream DNA storage methods based on synthesis and sequencing suffer from slow read/write speeds, hindering practical applications. To address this limitation, we propose an innovative hybridization-based DNA storage scheme. Through principle validation experiments, short-text read/write and error-correction tests, and high-density multi-format file storage experiments, we verify the feasibility of this approach. The proposed method exhibits high parallelism, rapid read speeds, and robust encryption capabilities, demonstrating promising application prospects in scenarios such as confidential data storage.

## Introduction

With the exponential growth of digital information driven by cloud computing, 5G communication, and artificial intelligence, traditional silicon-based storage media face severe bottlenecks due to the lagging supply of wafer-level silicon ^[1,2]^. DNA, as a natural genetic material, serves as an exceptional information carrier. By mapping digital data to nucleotide sequences, information can be stored and retrieved through DNA synthesis and sequencing. Theoretically, hundreds of grams of DNA could store all existing digital information^[2-4]^, with preservation times exceeding millennia—far surpassing silicon-based media ^[5]^.

Early attempts at DNA storage date back to the late 1980s ^[6]^. With advancements in second-generation sequencing, Church’s team achieved the storage of 658 KB of data in 2012 ^[3]^, marking a milestone in DNA storage research. Over the past decade, progress has been made in the following areas^[7-18]^:

Increasing Storage Capacity and Density: Methods include multi-symbol encoding with degenerate bases ^[19]^, high-density microarrays ^[20]^, expanded encodable sequences ^[21]^, and nested file addressing systems ^[22]^.

Enhancing Retrieval and Random Access: Techniques involve hybridization probes ^[23,24]^, primer-based amplification ^[25]^, structured data encoding^[7,8,24]^, DNA barcoding ^[12]^, and enzymatic approaches ^[13]^.

Improving Stability, Error Correction, and Encryption: Strategies include anhydrous/anaerobic preservation ^[26]^, comparative preservation studies ^[27-29]^, molecular bias analysis ^[30]^, fountain codes ^[8,31]^, forward error correction codes ^[32]^, HEDGES codes ^[33]^, Raptor codes ^[11]^, biologically constrained encoding ^[34]^, matrix-based encoding ^[35]^, STR encryption ^[36]^, and multi-encoding composite keys ^[37]^.

Advancing Synthesis/Sequencing Technologies: Innovations encompass CRISPR-based mutation coverage ^[18]^, photolithographic enzymatic synthesis ^[38,39]^, artificial chromosomes ^[40]^, parallelized synthesis platforms ^[20,41]^, portable random-access systems ^[9]^, integrated read/write systems ^[42]^, and self-contained storage architectures ^[43]^.

Most existing methods rely on sequential synthesis and sequencing, which remain time-consuming despite parallelization ^[20,41]^. Alternative approaches, such as nanopore-based secondary structure detection ^[44]^or methylation-based epigenetics ^[45]^, still depend on synthesis/sequencing. Here, we present a paradigm-shifting hybridization-based DNA storage technology ^[46-50]^, where binary data is encoded by the presence/absence of specific DNA strands on a microarray. Information is retrieved via parallel hybridization rather than sequencing, enabling rapid readout. Although this sacrifices some density, it achieves unprecedented read speeds and encryption potential, making it ideal for secure, time-sensitive applications.

## Results

### Principle Validation of 2^n^ State Discrimination Using n Encoding Strands

The encoding principle maps each binary bit to the presence (1) or absence (0) of a specific DNA strand. For n strands, 2^n^ states can be encoded. Using four encoding strands, we immobilized all 16 possible combinations (2□) on NHS-activated magnetic beads (Beaver Biosciences). Four fluorescent probes (Alexa 488, Cy3, Cy5, Cy7; Sangon Biotech), complementary to the strands, were hybridized. Fluorescence intensity analysis (Fig. 1) revealed significant differences (p < 0.001) between presence/absence states for each strand, confirming unambiguous discrimination of all 16 states.

**Figure 1.**
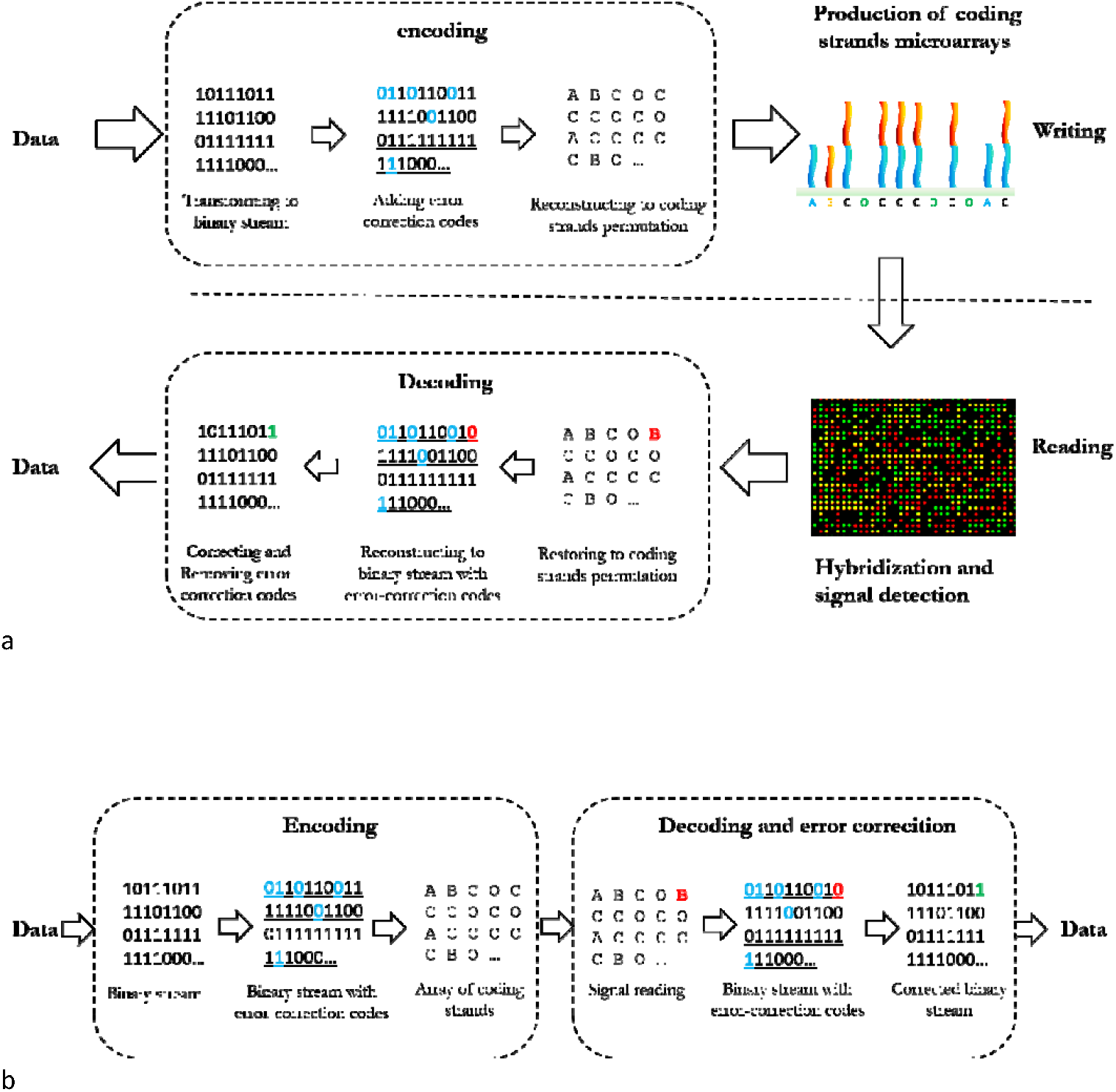
Schematic diagram of the principle of DNA hybridization storage (a) and error correction (b)

**Figure 2.**
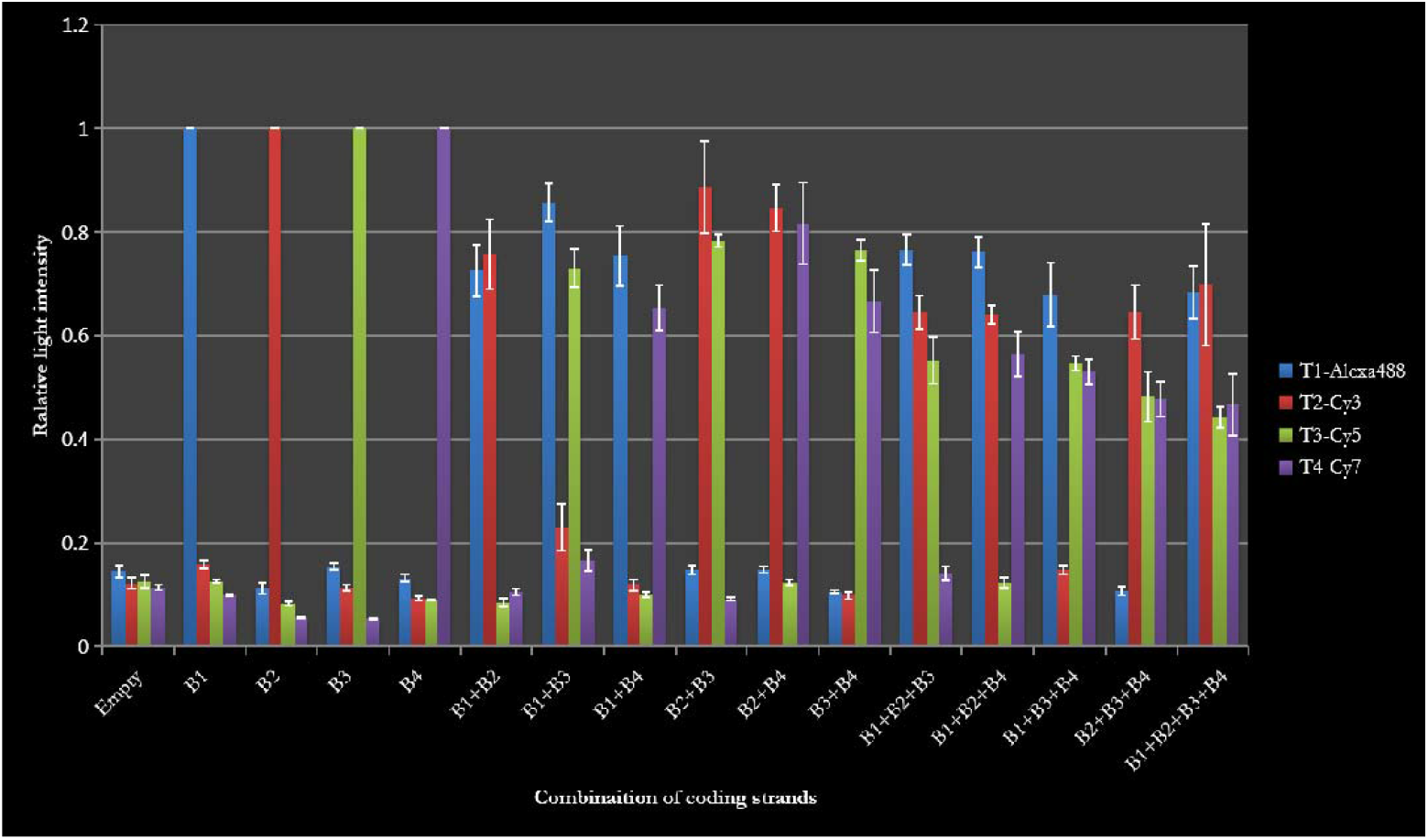
Experimental results of the principle verification of DNA hybridization storage When using four oligonucleotide coding strands, all 2^4=16 different combinatorial states can be effectively distinguished;

### Read/Write Testing of Chinese and English Short Texts

A dual-color encoding system (2 bits per data unit) was tested using a 744-bit Chinese poem (“Viewing the Lushan Waterfall”) and a 768-bit English quote (“Ask not what your country can do for you…”). Custom software converted texts to binary, mapped to a 26×20 microarray layout. Arrays were printed using an I-DOT dispenser (Sangon Biotech) on aldehyde-coated slides (Nexterion Slide AL, SCHOTT). Hybridization with Cy3/Cy5 probes (Sangon Biotech) and subsequent scanning (LuxScan-10K/A) yielded fluorescence signals decoded via AGScan software. Both texts were recovered with >99.9% accuracy, validating end-to-end functionality.

### High-Density Multi-Format File Storage

A myarray® custom microarray (Arbor Bio) stored 12.6–14.6 KB files (text, MP3, JPG, GIF). Read accuracy exceeded 99.98% per data unit. Text and audio files were flawlessly restored, while compressed images (JPG/GIF) exhibited minor artifacts (Fig. 3) due to error propagation in lossy formats. Error analysis revealed sparse, random bit errors (<0.02%), correctable via algorithms.

**Figure 3.**
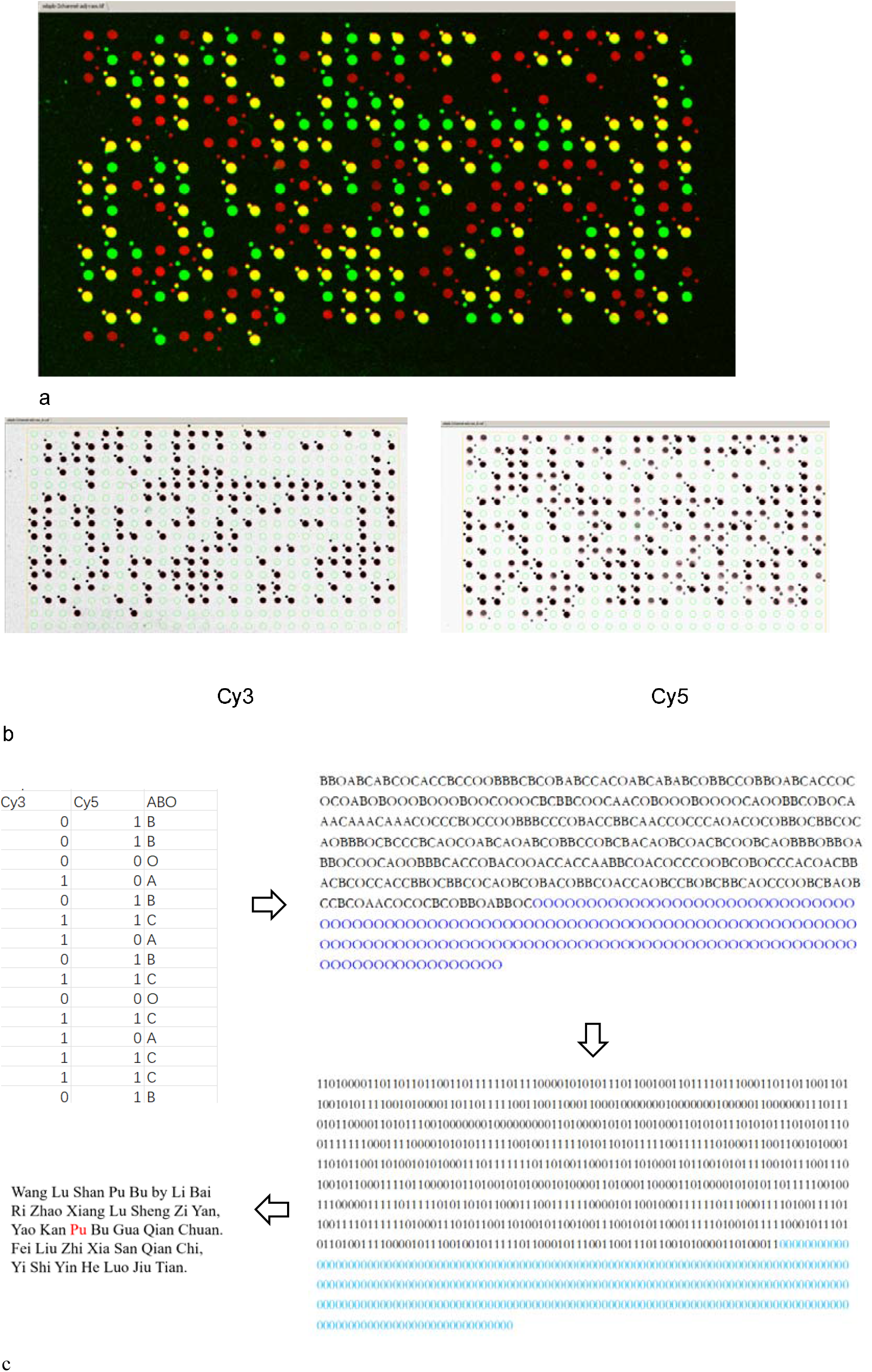
Experimental results of short text read/write in DNA hybridization storage

### Error Correction with Hamming Codes

A (16,11) Hamming code (k=11 data bits, r=5 parity bits) was implemented. With a raw error rate <0.02%, simulations predicted >93% full-file recovery probability (Fig. 4a). Empirical tests under suboptimal hybridization (e.g., contamination clusters) confirmed successful error correction (Fig. 4b).

**Figure 4.**
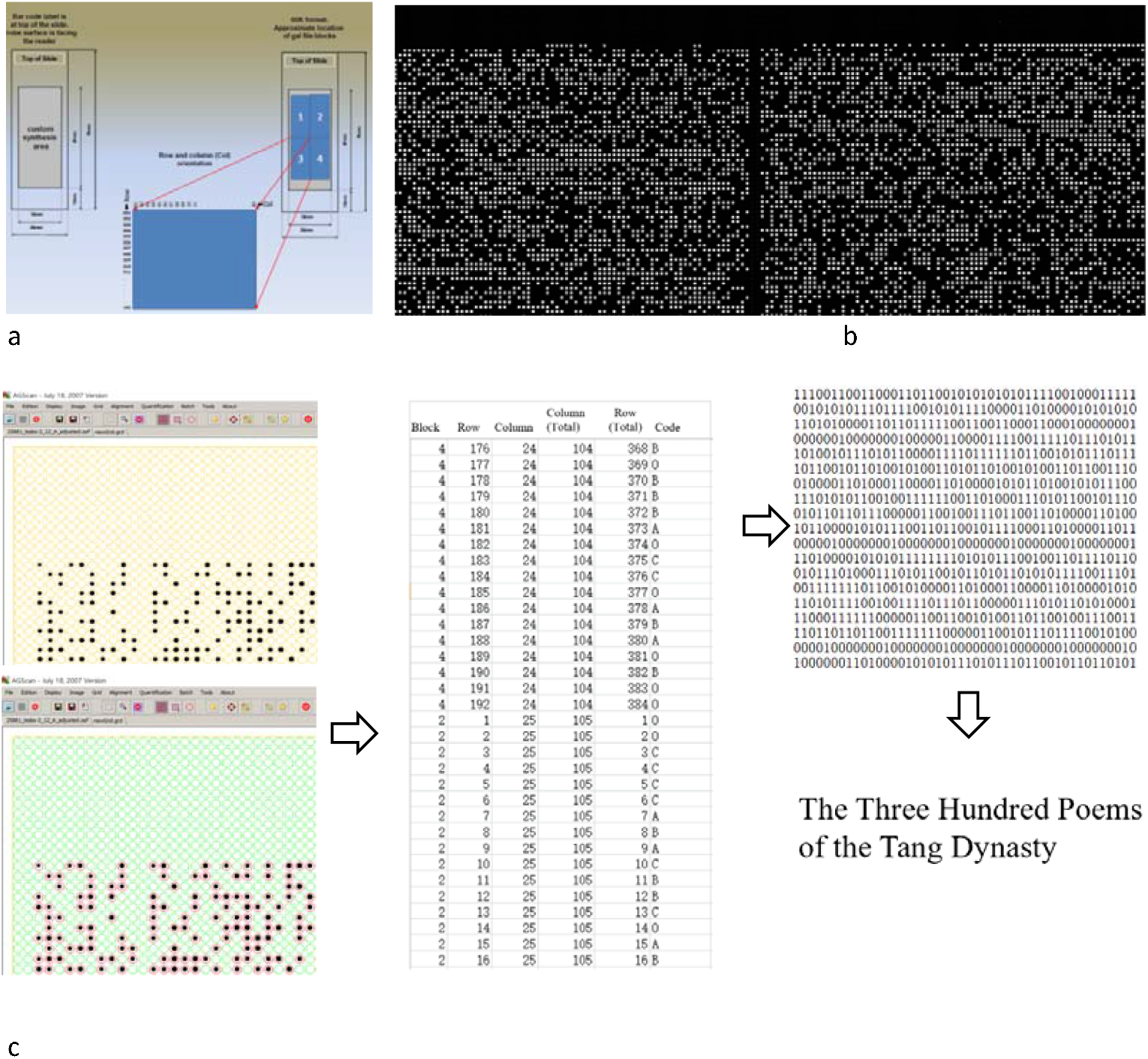
Experimental results of multi-format file access in DNA hybridization storage a. Specifications of the Myarray chip, b. Scanned image after hybridization, c. Signal processing and data restoration

### Long-Term Stability

Arrays stored at room temperature for 24 months exhibited no significant degradation in read accuracy (Fig. 5), consistent with DNA’s archival stability.

**Figure 5.**
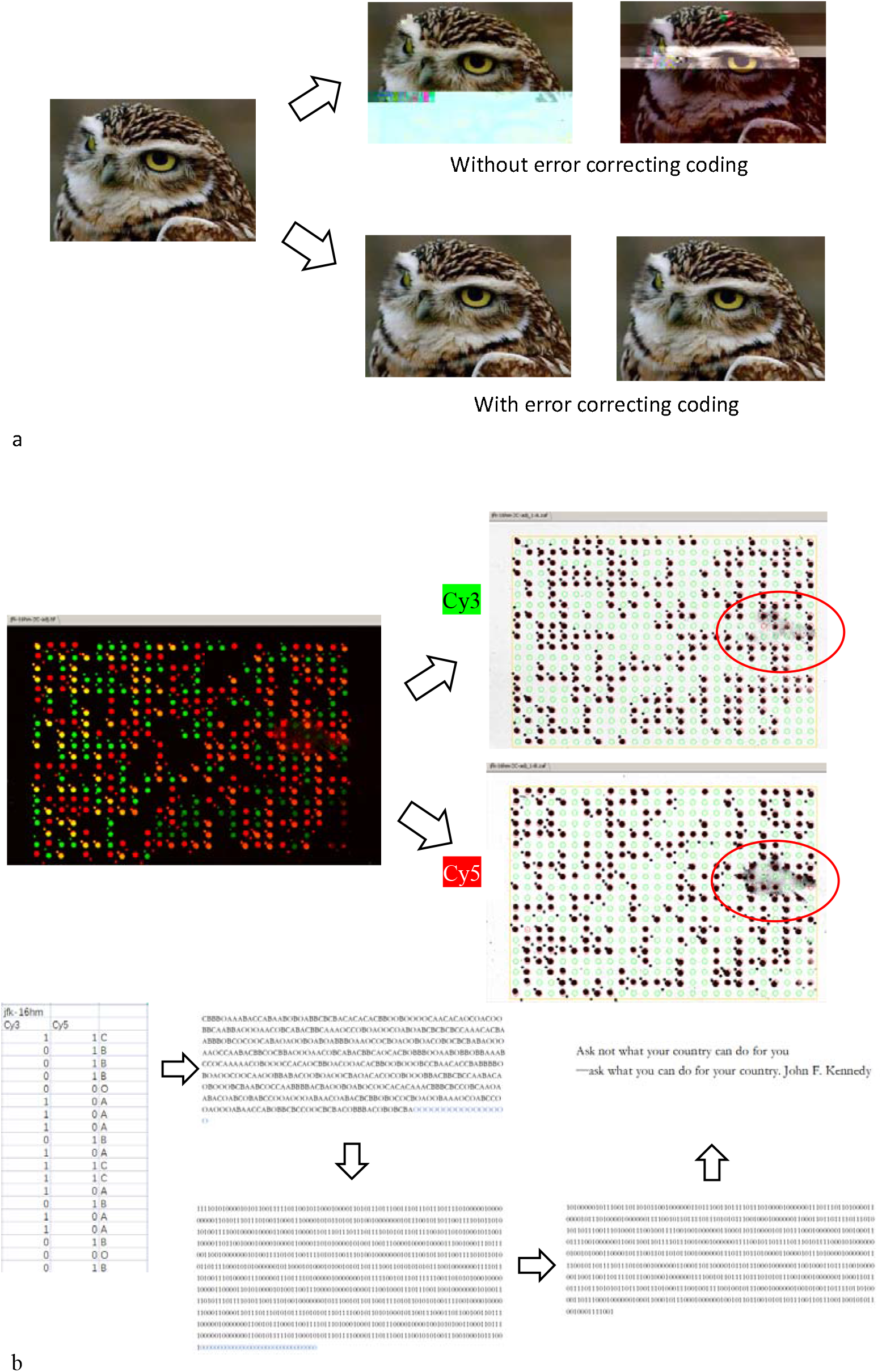
Experimental results of error correction in DNA hybridization storage a. Simulation experiment of error correction on the Myarray chip, b. Actual effect of short text error correction

**Figure 6.**
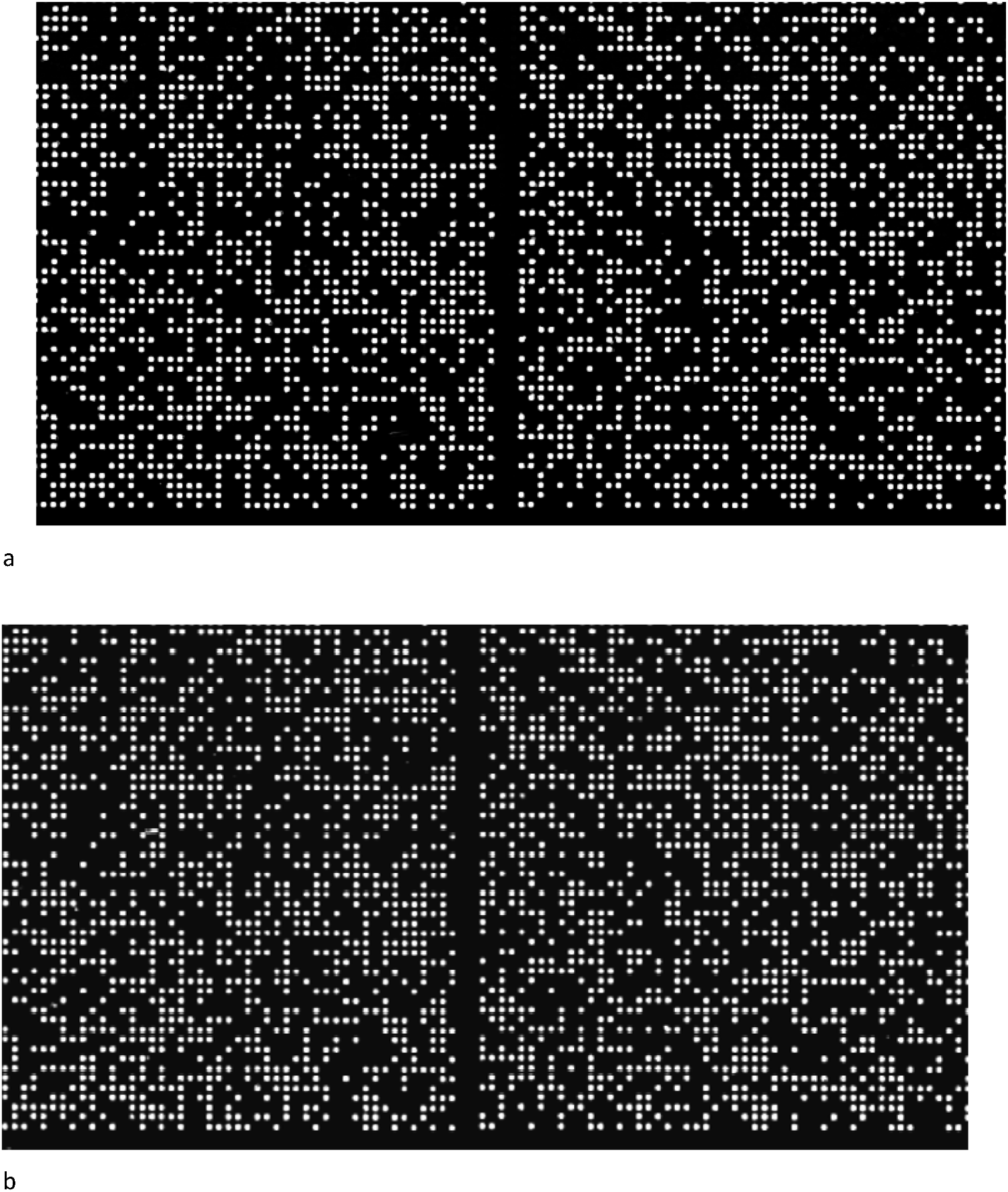
Results of the aging test for DNA hybridization storage. a. Scanned image of a chip hybridization after 1 month from production, b. Scanned image of another chip hybridization after 24 months from production for the same batch of products

### Discussion

Our hybridization-based DNA storage achieves rapid, parallelized reads while maintaining high accuracy and encryption potential. Although density lags behind sequence-based methods, its speed advantage suits secure, small-to-medium data storage. Current write speeds rely on conventional microarray printing; future work will explore prefabricated oligonucleotide linking to accelerate writing.

## Methods

### Principle Validation

Four amino-modified DNA strands (Sangon Biotech) were immobilized on NHS-activated magnetic beads. Hybridization with fluorescent probes (Sangon Biotech) was analyzed using a SpectraMax M2 microplate reader (Molecular Devices).

### Text Storage

Texts were converted to binary, mapped to a 26×20 microarray. Arrays were printed using an I-DOT dispenser with amino-modified DNA oligos (Sangon Biotech) on aldehyde slides (Nexterion Slide AL, SCHOTT). Hybridization and scanning protocols matched the validation experiments.

### Multi-Format Storage

Files were encoded for Myarray® microarrays (Arbor Bio), hybridized with Cy3/Cy5 probes, and scanned using an Agilent DNA Microarray Scanner 2100.

### Error Correction

Hamming codes (k=11, r=5) were applied during encoding/decoding.

## Acknowledgments

This work was supported by the National Key R&D Program of China (2020YFA0907000, Di Liu), and the National Natural Science Foundation of China (32270528, Yifei Zhang). We thank Yaowen Zhang (Huazhong University of Science and Technology), Bo Feng (Xiangtan University) and Dynegene Technologies for technical assistance.

